# Structural insights into dual-antagonize mechanism of AB928 on adenosine A_2_ receptors

**DOI:** 10.1101/2023.07.01.547314

**Authors:** Yuan Weng, Xinyu Yang, Qiansen Zhang, Ying Chen, Yueming Xu, Chenyu Zhu, Qiong Xie, Yonghui Wang, Huaiyu Yang, Mingyao Liu, Weiqiang Lu, Gaojie Song

**Affiliations:** Shanghai Key Laboratory of Regulatory Biology, Institute of Biomedical Sciences and School of Life Sciences, East China Normal University, 500 Dongchuan Road, Shanghai 200241, China; Department of Medicinal Chemistry, School of Pharmacy, Fudan University, 826 Zhangheng Road, Shanghai 201203, China

## Abstract

The adenosine subfamily G protein-coupled receptors A_2A_R and A_2B_R were identified as promising candidates for cancer immunotherapy within recent years. One of the A_2A_R/A_2B_R dual antagonist, AB928, has progressed to phase II clinic trial for the treatment of rectal cancer. However, the precise mechanism underlying its dual-antagonistic properties remains elusive. Herein, we report crystal structures of A_2A_R in complex with AB928 and a selective A_2A_R antagonist, 2-118. The structures reveal a common binding mode on A_2A_R, wherein the ligands establish extensive interactions with residues from both the orthosteric pocket and the secondary pocket. Conversely, the cAMP assay together with molecular dynamics simulations conducted on both A_2A_R and A_2B_R indicate that the ligands adopt distinct binding modes on A_2B_R. Detailed analysis of their chemical structures suggests that AB928 can readily adapt to the A_2B_R pocket, while 2-118 cannot due to its intrinsic differences. This disparity potentially accounts for their divergent inhibitory efficacies between A_2B_R and A_2A_R. The findings from this study can serve as valuable structural templates for future development of selective or dual inhibitors targeting A_2A_R/A_2B_R in the context of cancer therapy.

## Introduction

Cancer immunotherapy has emerged as a potent strategy in the fight against neoplastic diseases, drawing significant attention from both academia and industry. One of the most promising approaches in this field involves the development of inhibitors for immune checkpoints, such as programmed cell death protein 1 (PD-1), programmed death-ligand 1 (PD-L1) and cytotoxic T-lymphocyte-associated protein 4 (CTLA-4) (Marin-Acevedo et al., 2018). These inhibitors have demonstrated significant success in treating related cancers. Recently, the CD39/CD73/adenosine axis has been identified as a crucial factor in suppressing immune responses and promoting tumor growth. This axis works by hydrolyzing extracellular ATP to AMP using CD39, and then further hydrolyzing AMP to adenosine with CD73. Adenosine then activates A_2A_R and A_2B_R receptors found on tumor cells, leading to enhanced tumor growth and proliferation. Consequently, inhibiting either hydrolase or antagonizing the adenosine receptors have been shown to be effective anti-tumor strategies (Saini et al., 2022; Yu et al., 2020).

The adenosine receptor family includes four members, A_1_R, A_2A_R, A_2B_R, and A_3_R, which can sense adenosine (Fredholm et al., 2001). Among these, A_2A_R and A_2B_R activate adenylyl cyclase (AC) and produce cyclic AMP (cAMP) by recruiting downstream stimulatory G protein (G_s_). In contrast, A_1_R and A_3_R trigger the opposite function by recruiting inhibitory G protein (G_i_). Although A_2B_R is less sensitive to adenosine than A_2A_R, it is activated along with A_2A_R in the tumor microenvironment (TME), where the concentration of adenosine is much higher than in normal tissue (Borea et al., 2018). Elevated intracellular cAMP in immune cells can lead to anergy of several immune cells, including natural killer cells, dendritic cells, and T cells (Beavis et al., 2015; Dziedzic et al., 2021; Hofer et al., 2021). Therefore, dual-antagonism of both A_2A_R and A_2B_R has been shown to be a promising strategy compared to selective antagonism.

Several small molecular antagonists against individual A_2A_R or A_2B_R have entered clinical trials to treat different cancers, such as CPI-444 (Iacovelli et al., 2022), AZD4635 (Lim et al., 2022), and PBF-509 (Chiappori et al., 2022) against A_2A_R, and PBF-1129 (Evans et al., 2023) and TT-4 (Pastore et al., 2021) against A_2B_R. Dual-antagonists, such as AB928 (Seitz et al., 2019) and M1069 (Zaynagetdinov et al., 2022), are also under clinical evaluation for the treatment of rectal cancer or solid tumors. The A_2A_R/A_2B_R dual-antagonist AB928 has been reported to outperform its competitors in preclinical tests (Walters et al., 2017), suggesting potential advantages for dual-antagonism. While the binding modes of ∼40 antagonists have been determined experimentally, there is no complex structure available for dual-antagonists like AB928, hence the molecular mechanism for dual-antagonism remains elusive. Here, we present high-resolution structures of A_2A_R in complex with the dual-antagonist AB928 and a close analogue of AB928 that selectively antagonizes A_2A_R. Our structures, together with molecular dynamics simulations and cell-based assay, reveal the mechanism for dual-antagonism and the potentially unique binding features on A_2B_R by these ligands.

## Results

### Crystal structures

To elucidate the molecular mechanism underlying the dual-antagonism of AB928, our investigation commenced with the crystallization of A_2A_R in complex with AB928 (3-[2-amino-6-[1-[[6-(2-hydroxypropan-2-yl)pyridin-2-yl]methyl]triazol-4-yl]pyrimidin-4-yl]-2-methylbenzonitrile). The crystallization construct of A_2A_R resembled that of the first inactive A_2A_R–ZM241385 structure (Jaakola et al., 2008), featuring a T4 lysozyme at intracellular loop 3 (ICL3). The A_2A_R–AB928 complex shared comparable thermal-stability to the A_2A_R–ZM241385 complex and crystallized into an identical lattice (Figure S1). The A_2A_R–AB928 complex crystals diffracted to 2.37 Å, permitting unambiguous modelling of the ligand (Figure 1A). The solved complex structure revealed that the AB928 not only inserted deep within the orthosteric pocket but also extended into the so-called secondary pocket composed of transmembrane helices 1, 2, and 7 (TM1/2/7) (Figure 1B) (Chen et al., 2022). Most strikingly, the methylbenzonitrile moiety at the head of AB928 formed hydrophobic contacts with residues V84^3.32^, L85^3.33^, M177^5.38^, W246^6.48^ and H250^6.52^, while its cyano group established a hydrogen bond with T88^3.36^. The central 2-aminopyrimidine moiety engaged in a π-π stacking interaction with the conserved F168 at extracellular loop 2 (ECL2) and formed hydrogen bonds with E169^ECL2^ and N253^6.55^, reminiscent of other A_2A_R antagonists or agonists of A_2A_R. The adjacent triazole moiety of AB928 exhibited minimal contact with the receptor, except for indirect hydrogen bonds with S277^7.42^ and H278^7.43^ via a water network (Figure 1B). The position of this triazole moiety is likely stabilized by the pyridine ring at the tail of AB928, which makes extensive non-polar interactions with residues Y9^1.35^, A63^2.61^, S67^2.65^, Y271^7.36^ and I274^7.39^ in the secondary pocket. Furthermore, a 2-hydroxyisopropyl moiety attached to the 2-position of the pyridine ring interacted with L267^7.32^, M270^7.35^ and Y271^7.36^, with its hydroxyl group exposed to the solvent environment at the edge of the orthosteric pocket (Figure 1B).

**Figure 1.**
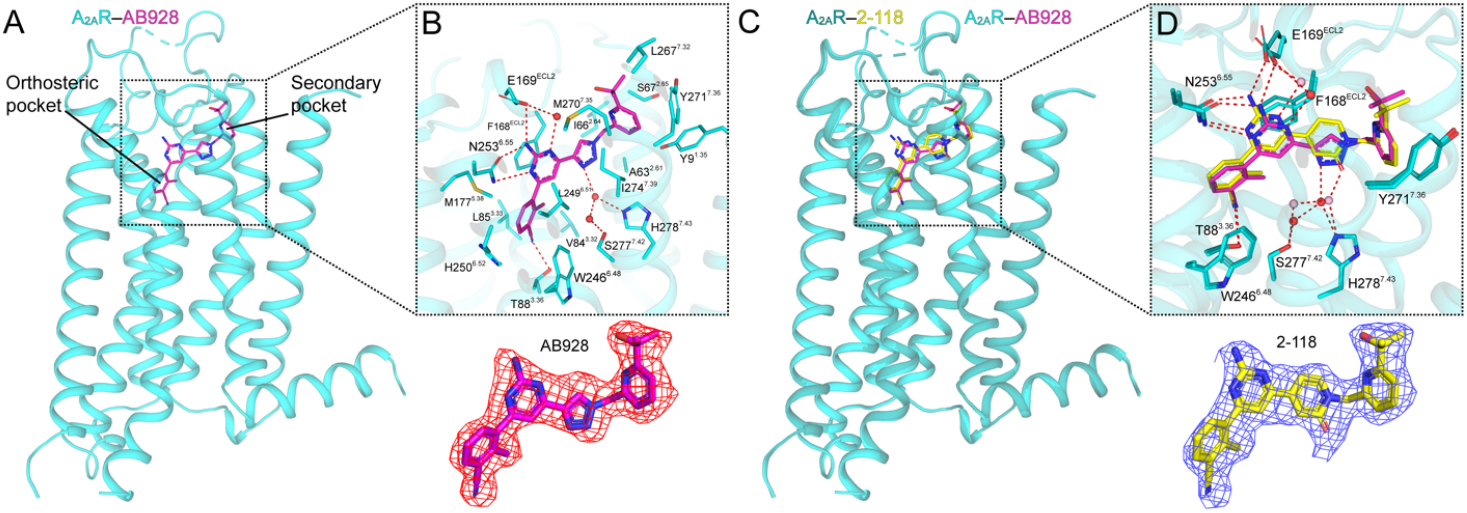
The binding modes of AB928 and 2-118 on A_2A_R. A, overall structure of A_2A_R (cartoon, cyan) binding AB928 (sticks, magenta). B, detailed binding pocket of AB928, sidechains of the interacting residues are shown as sticks, water molecules are shown as red spheres, hydrogen bonds are shown as red dashes. Density of AB928 is shown below as red meshes. C, overall structure of A_2A_R (cartoon, teal) binding 2-118 (sticks, yellow) superimposed with that of A_2A_R– AB928. D, detailed binding pocket of 2-118 superimposed with that of AB928, sidechains of the interacting residues are shown as sticks, water molecules are shown as pink spheres. Density of 2-118 within the pocket is shown below as blue meshes.

We recently reported structure-activity relationship study of a series of A_2A_R antagonists (Zhu et al., 2023), in which we described a close analogue of AB928, compound 40 (hereinafter referred to as 2-118). 2-118 (3-[2-amino-6-[1-[[6-(2-hydroxypropan-2-yl)pyridin-2-yl]methyl]-2-oxo-1,2-dihydropyridin-4-yl]pyrimidin-4-yl]-2-methylbenzonitrile) has a structure almost identical to that of AB928, except for the replacement of the triazole moiety in AB928 with a pyridinone ring. However, compared to AB928, although 2-118 shows comparable antagonism towards A_2A_R, its activity on A_2B_R is greatly compromised (Zhu et al., 2023). To further explore the distinctions between these two ligands, we resolved the A_2A_R structure bound to 2-118 using a similar approach (Figure 1C).

The binding model of 2-118 closely resembles that of AB928, with the methylbenzonitrile and pyridine rings in 2-118 aligning perfectly with the corresponding head and tail moieties of AB928 (Figure 1D). The pyridinone ring of 2-118 partially occupies the position held by the triazole ring in AB928, with its carbonyl group oriented into the pocket, forming indirect hydrogen bonds with residues in TM7 (Figure 1D). The water molecules beneath the pyridinone/triazole rings are conserved in both structures, suggesting their involvement in the interactions between the antagonists and A_2A_R. Notably, the altered connection introduced by the pyridinone moiety in 2-118 induces a ∼1 Å movement of its pyrimidine moiety, resulting in a similar shift in the side chains of F168^ECL2^, E169^ECL2^, and N253^6.55^. Nevertheless, despite these minor movements within the pocket, the overall structures of the 2-118- and AB928-bound A_2A_Rs are highly similar to each other (Cα RMSD 0.18 Å) and exhibit an essentially identical conformation with previous inactive A_2A_R structures (Figure 2).

**Figure 2.**
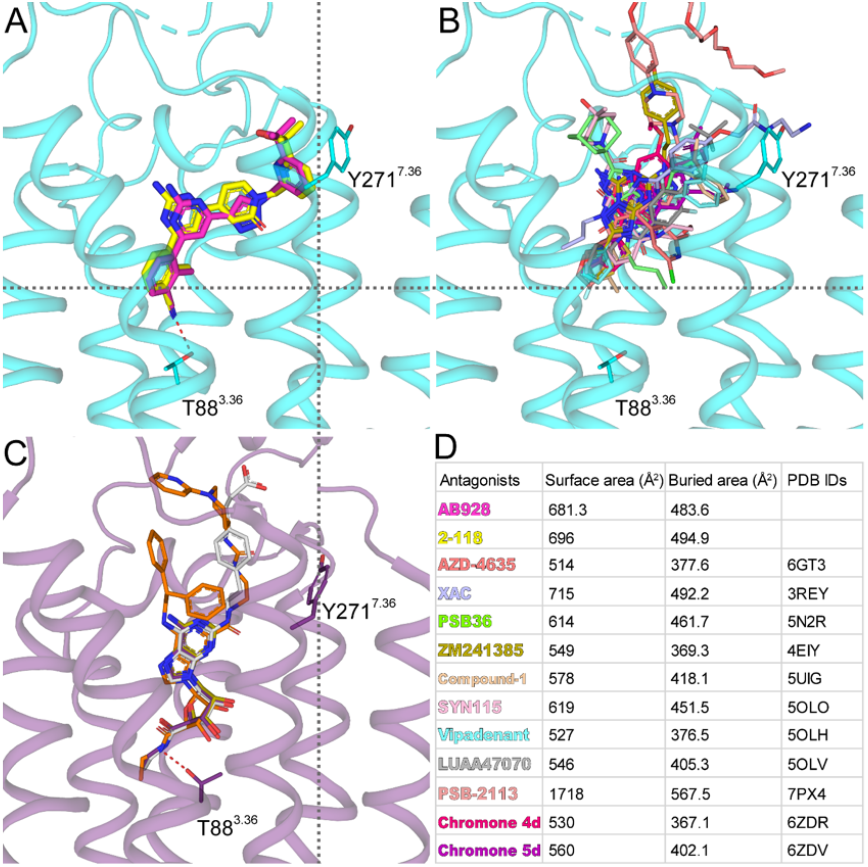
Comparison of the AB928/2-118-bound A_2A_R structures with other antagonist or agonist-bound A_2A_R structures. A-C, sideview of the pockets of A_2A_R binding AB928/2-118 (A), other representative antagonists (B), and agonists (C). Locations of landmark residues T88^3.36^ and Y271^7.36^ are shown as sticks. In (C) representative antagonist-bound structures are: AZD-4635 (6GT3, deep salmon), XAC (3REY, light blue), PSB36 (5N2R, green), ZM241385 (4EIY, light olive), Compound-1 (5UIG, wheat), SYN115 (5OLO, pink), Vipadenant (5OLH, cyan), LUAA47070 (5OLV, deep grey), PSB-2113 (7PX4, salmon), Chromone 4d (6ZDR, hot pink) and Chromone 5d (6ZDV, purple). In (D) structures of CGS21680 (4UHR, light grey), UK-432097 (3QAK, orange), adenosine (2YDO, deep olive) binding A_2A_R are superimposed with that of NECA (PDB ID 2YDV, deep purple). D, comparison of the surface areas buried by different antagonists.

The AB928- or 2-118-bound A_2A_R structures provide insights into why AB928 outperforms other antagonists (Walters et al., 2017) (Figure 2, S2). Comparing the locations of two landmark residues, T88^3.36^ and Y271^7.36^, revealed that AB928 or 2-118 occupies the broadest and deepest position in the pocket of A_2A_R and makes contacts with all helices except TM4 (Figure 2A and B). In contrast, adenosine derivatives, which are agonists for adenosine receptors, occupy only a partial space in the horizontal direction but insert even deeper in the vertical direction (Figure 2C). The association of agonists with deep positions like 3.40 and 6.48 was suggested to be key in triggering the conformational change for receptor activation (Zhou et al., 2019). Figure 2D showed that the surface areas buried by AB928/2-118 are among the highest of all buried surfaces of different antagonists. Therefore, it is speculated that AB928/2-118 may have made the best use of the space within the pockets to antagonize receptor activation.

### Determinants in the secondary pocket

Previous studies have highlighted the importance of specific residues within the orthosteric pocket for A_2A_R function (Borodovsky et al., 2020; Doré et al., 2011; Liu et al., 2012; Sun et al., 2017), including F168^ECL2^, N253^6.55^ and W246^6.48^. However, fewer studies have explored the secondary pocket. AB928/2-118 packed tightly against Y271^7.36^ in the secondary pocket (Figure 3A); however, this position is not conserved in A_2B_R, where an equivalent position is occupied by a glutamine. To validate the function of Y271 in AB928/2-118 recognition we introduced a Y271^7.36^N mutation in A_2A_R and measured the inhibitory potency of AB928/2-118. The results demonstrated that the mutation indeed decreased the potency of AB928 and 2-118 by ∼20- and ∼30-fold, respectively (Figure 3B and C). This outcome is aligned with the observation that Y271^7.36^ was displaced away from the core of the helices by 1.6-4.3 Å compared to other A_2A_R structures (Figure 3D), suggesting an essential role for Y271 in A_2A_R’s recognition of AB928 and 2-118. In each structure, the side-chain of Y9 points towards the base of the pyridine ring (Figure 3A), while its hydroxyl group at the tip appears to be incompatible with the hydrophobic pyridine ring in the ligand. Consistently, removal of the hydroxyl group by the Y9^1.35^F mutation slightly increased the potency of AB928 (1.8-fold) and 2-118 (2.7-fold) (Figure 3B and C).

**Figure 3.**
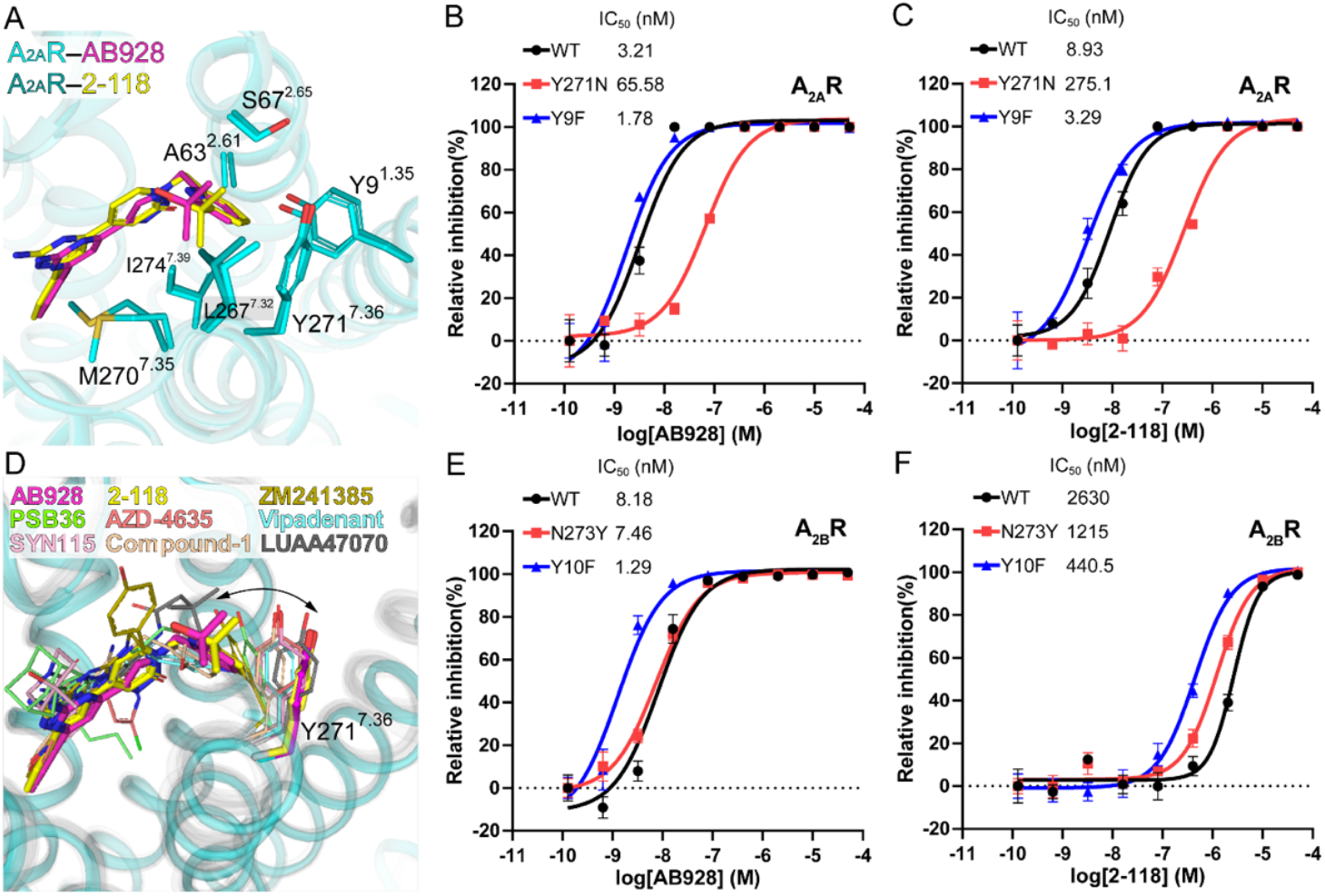
Signaling profiles of key residues in the secondary pocket of A_2A_R and A_2B_R. A, Close-up view of the secondary pocket. Color codes are same as Figure 1. Secondary pocket residues are shown as sticks. B-C, mutagenesis analysis of key residues in the secondary pocket of A_2A_R on the potency of AB928 (B) and 2-118 (C). D. Close-up view of the superimposed secondary pocket of different antagonist-bound A_2A_R structures. Color codes are shown on the top. Y271^7.36^ in each structure is shown as sticks and colored the same as the antagonist. AB928-bound A_2A_R is colored cyan and all other A_2A_R are colored grey. dynamics of Y271^7.36^ is marked by a double-headed arrow. E-F, mutagenesis analysis of key residues in the secondary pocket of A_2B_R on the potency of AB928 (E) and 2-118 (F). Data are shown as means ± SEM from at least 3 independent experiments.

Our cAMP assay confirmed that AB928 can antagonize A_2B_R to a similar single-digit nanomolar level as A_2A_R (Figure 3B and E), in contrast to the ∼20-fold reduction by the Y271^7.36^N mutant of A_2A_R (Figure 3C and F). Interestingly, the corresponding N273^7.36^Y mutation on A_2B_R did not significantly enhance the potency (IC_50_=7.46 nM), unlike the ∼6-fold increase in potency observed with the Y10^1.35^F mutation on A_2B_R. These findings suggested that although AB928 may bind to the pocket of A_2B_R in a similar manner, the pyridine ring likely adopts a slightly different orientation that makes little contact with position 7.36. This analysis is also in line with the incompatible feature between the hydrophobic pyridine ring and the polar N273^7.36^ residue. Meanwhile, compared to AB928, the potency of 2-118 decreased by three orders of magnitude on A_2B_R (IC_50_=2.63 uM), and the N273^7.36^Y and Y10^1.35^F mutations on A_2B_R only partially rescued the potency by ∼2- and ∼6-fold, respectively (Figure 3E and F). Together, these results indicated that, although the AB928 and 2-118 adopt similar binding poses in A_2A_R, they probably adopt different binding poses in A_2B_R, resulting in distinct pharmacological effects. Moreover, the determinate residue(s) controlling the binding capacity and potency of these two ligands differ between A_2B_R and A_2A_R.

### MD simulations on AB928/2-118-bound A_2A_R and A_2B_R

To further gain insights into the effects of AB928/2-118, we performed molecular dynamics (MD) simulations on both adenosine receptors. Firstly, we conducted side-by-side MD simulation for the A_2A_R–AB928 and A_2A_R–2-118 structures, each lasting ∼ 500 ns. The root mean square deviation (RMSD) of ∼1 Å for each A_2A_R ligand suggested a stable orientation for the ligands, and the receptor remained in an inactive conformation (Figure 4A and B). Quantitatively, T88 preserved hydrogen-bond interactions with AB928 and 2-118 to average percentages of 91.1% and 90.9%, respectively (Figure 4C and D). Despite minor fluctuations, the Y271^7.36^ was mostly stabilized in its original position due to hydrophobic interactions with the pyridine ring of each ligand. These features aligned well with their strong antagonistic activities.

**Figure 4.**
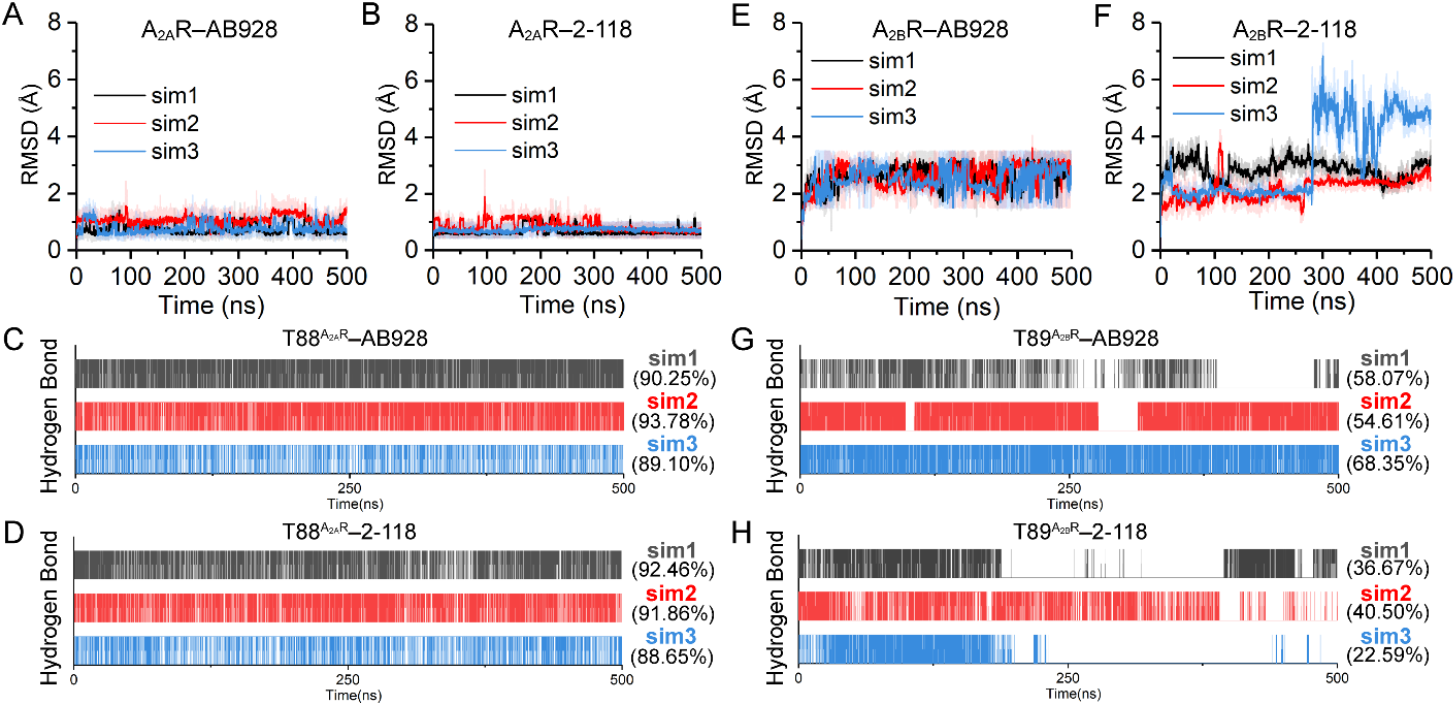
MD simulations of the A_2A_R/A_2B_R in complex with AB928/2-118. A-B, RMSD of AB928 (A) or 2-118 (B) from 500 ns MD simulation with A_2A_R (crystal structures as the starting models). C-D, statistics of hydrogen bond interaction between T88 of A_2A_R and AB928 (C) and 2-118 (D). E-F, RMSD of AB928 (E) or 2-118 (F) from 500 ns MD simulation with modelled A_2B_R. G-H, statistics of hydrogen bond interaction between T89 of A_2B_R and AB928 (G) and 2-118 (H). Experiments are performed with triple trajectories, abbreviated with sim1-3 respectively.

The starting model of A_2B_R was built based on the A_2A_R structures, and the ligands were docked in a similar manner. Nevertheless, the subsequent MD simulations of modelled AB928 or 2-118 in A_2B_R exhibited larger fluctuations for both ligands (Figure 4E and F). Consistently, the T89^3.36^– AB928 and T89^3.36^–2-118 hydrogen bonds were only partially maintained in the A_2B_R models (Figure 4G and H). In contrast to the cyano group that inserted deep into the orthosteric pocket, the pyridine ring displayed greater dynamic behavior during the simulation, and in later stage, it even moved out of the secondary pocket (Figure 4F, simulation 3). It is worth to mention that AB928 performed relatively better than 2-118 in the simulations, as indicated by lower RMSD values and higher percentages of hydrogen bonding (average 60% vs 33.2%).

### AB928/2-118 adopt distinct binding modes in A_2B_R

The relatively dynamic feature of AB928 on A_2B_R over A_2A_R seems contradictory to the pharmacological data, which showed a similar level of potency on both receptors. However, the MD simulations have nevertheless provided potential hints into the docking model of AB928 in the A_2B_R pocket (Figure 3 and 4). In many snapshots, the ligand exhibited a slight clockwise rotation from a top view, and the pyridine moiety at the tail of AB928 twisted towards ECL3, forming interactions with two residues, K267^ECL3^ and K269^7.32^ (Figure S3). These two basic residues in A_2B_R are not conserved within the adenosine receptor subfamily and correspond to A_2A_R residues A265^ECL3^ and L267^7.32^, respectively. Hence, to explore the role of these residues we individually mutated them to alanine and test their functional consequences. The cAMP assay results on A_2B_R showed that the K267^ECL3^A and K269^7.32^A mutations reduced the potency of AB928 by 3.8- and 4.2-fold, respectively (Figure 5A). Remarkably, the same mutations conversely increased the potency of 2-118 on A_2B_R by 4-5-fold (Figure 5B). These results suggest that the K267^ECL3^ and K269^7.32^ are indeed involved in the interaction with AB928, potentially compensating for the absence of a bulky side-chain at position 7.36 in A_2B_R. On the contrary, the bulky side-chains on K267^ECL3^ and K269^7.32^ may impede the docking of 2-118, thus removal of these side-chains are beneficial for the inhibitory function of 2-118. Hence, these results further suggested that although the AB928 and 2-118 adopt similar binding poses in A_2A_R, they may adopt different binding poses in A_2B_R according to the simulations and mutagenesis results.

**Figure 5.**
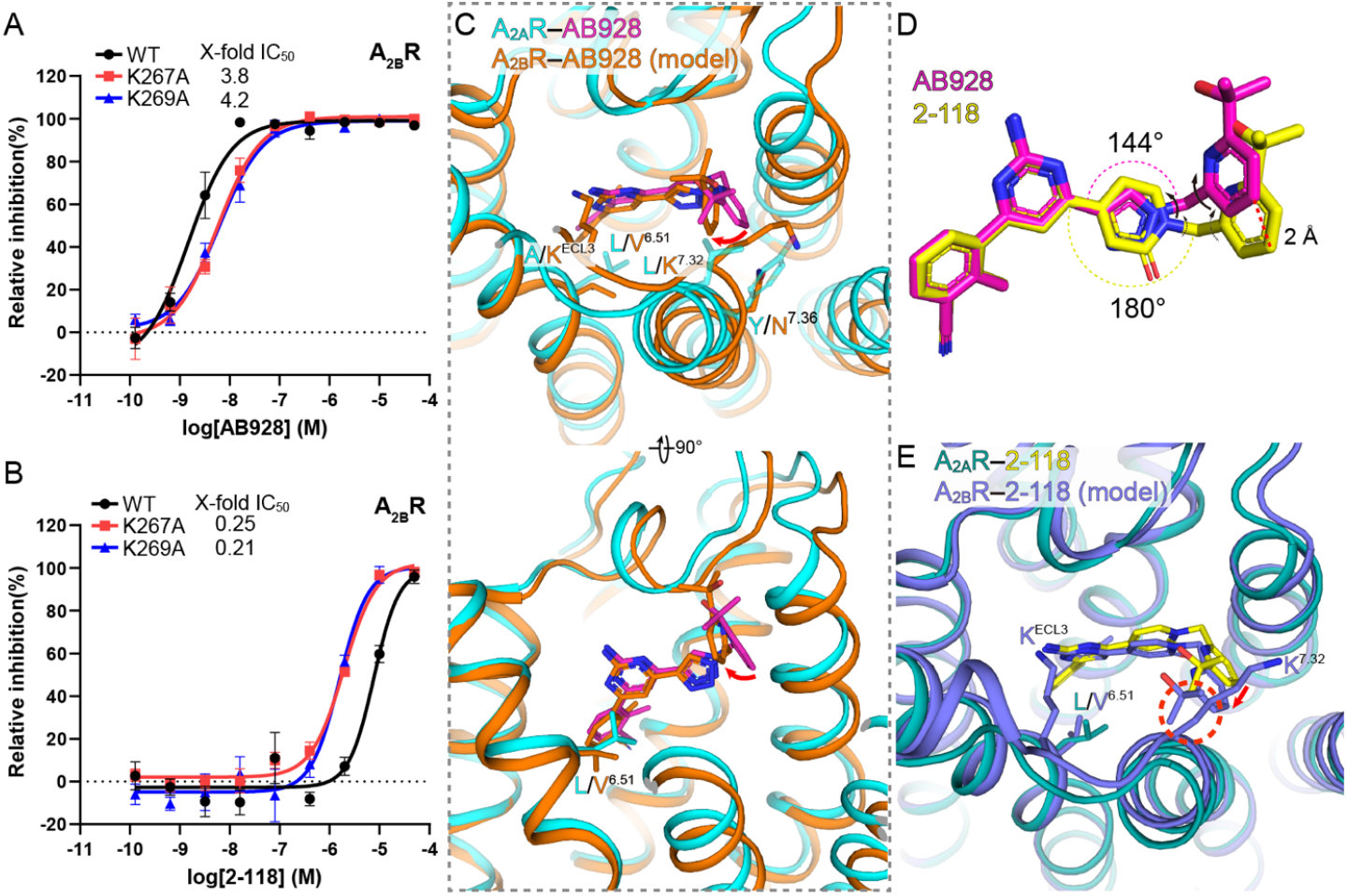
The AB928/2-118 adopt diverse binding poses on A_2B_R. A-B, mutagenesis analysis of residues K267^ECL3^ and K269^7.32^ of A_2B_R on the potency of AB928 (A) and 2-118 (B). Data are shown as means ± SEM from at least 3 independent experiments. C. superposition of the plausible model of A_2B_R–AB928 (based on a snapshot during simulations) onto the A_2A_R–AB928 crystal structure. Key sidechains are shown as sticks, rotation of the ligand is marked by a bent red arrow. D. Comparison of the chemical structures of AB928 and 2-118. These compounds are superimposed on the 2-aminopyrimidine moiety. The arrows indicate the groups can rotate around the single bonds, and the thin arrows on 2-118 mean restricted rotation. E. Extracellular view of a similar binding model of 2-118 on A_2B_R as in C. Rotation of the ligand is marked by a straight red arrow. Potential steric clash is marked by a dashed red circle.

Previous references have identified a key position (6.51) within the orthosteric pocket (Chen et al., 2022; Wang et al., 2021), which is a leucine in A_2A_R but occupied by a smaller residue, V250^6.51^, in A_2B_R. The active structures of A_2A_R/A_2B_R bound to 5’-N-ethylcarboxamidoadenosine (NECA) revealed that the ribose moiety of NECA undergoes a rotation and moves toward the V250^6.51^ in A_2B_R to establish hydrophobic contacts (Figure S4A). Combining this knowledge with our results strongly supports a specific snapshot, in which the methylbenzonitrile moiety of AB928 adopts a similar position as in A_2A_R and hydrogen bonds to the T89^3.36^, while the pyrimidine moiety rotates and establishes similar contacts with V250^6.51^ in A_2B_R (Figure S4B). Fine-tuning of the pyrimidine moiety further results in a slant upward movement of the pyridine ring by 2.5 Å, with one of the methyl groups at the tail being flanked by the bulky side-chains of K267^ECL3^ and K269^7.32^ (Figure 5C).

In contrast to a plausible model for the AB928–A_2B_R complex and the high potency of AB928, 2-118 exhibits significantly weaker inhibition, suggesting a similar binding model may not applicable to 2-118. The only difference between 2-118 and AB928 lies in the pyridinone ring, with the 2-aminopyrimidine and pyridine moieties connected to the para-positions of the pyridinone ring in 2-118. In AB928, the same moieties are connected to the triazole ring through the 4’ and 1’ positions, resulting in an angle of ∼144° between the 2-aminopyrimidine and pyridine moieties (Figure 5D). This 36° angle difference leads to a lift of the pyridine tail by ∼2 Å in AB928 compared to 2-118 when their 2-aminopyrimidine moieties are superimposed (Figure 5D). The lower position of the pyridine ring in 2-118 may cause steric incompatible between the pyridine moiety and the extracellular tip of TM7 when the pyrimidine moiety rotates towards TM6 to accommodate the shorter side-chain of V250^6.51^ (Figure 5E). The mutagenesis data showing that the K267^ECL3^A and K269^7.32^A significantly improve the efficacy of 2-118 are supportive of this hypothesis. Nevertheless, we cannot rule out that other residues from ECLs of A_2B_R may also played critical roles in the recognition of AB928 and 2-118. The detailed mechanism may rely on determination of the high-resolution inactive structure of A_2B_R in complex with AB928 and 2-118.

## Discussion

Here we determined crystal structures of A_2A_R in complex with the A_2A_R/A_2B_R dual antagonist AB928 and a A_2A_R-selective antagonist 2-118. The structures revealed a common binding mode on A_2A_R in which the ligands form extensive interactions with residues from the orthosteric and secondary pockets. The complex structures can explain many pharmacological data on AB928 or 2-118 derivatives. For example, the hydrogen bond contributed by the cyano group explained why the methylbenzonitrile moiety is better than a furan moiety in the corresponding position of many antagonists, and the interactions contributed by the 2-hydroxyisopropyl moiety account for why it is superior to other substituents, and why the substitution should be located at the 2’ position (Zhu et al., 2023).

Notably, an unprecedent hydrogen bond interaction occurs between T88 of A_2A_R and the cyano group of AB928/2-118. Such a polar contact is commonly observed between A_2A_R and its agonists, such as NECA, but has never been seen in previous antagonist-bound A_2A_R structures. Since T88 is conserved throughout the adenosine receptor subfamily, this feature may be utilized in future design of antagonists for adenosine receptors. Additionally, the insertion of the pyridine ring of AB928 into the secondary pocket may further explain why AB928 outperforms other antagonists, as extending the ligand from orthosteric pocket to the secondary pocket also displaces several water molecules within the ligand binding pocket. In comparison to high-resolution structures of A_2A_R in complex with ZM241385 (PDB: 4EIY) (Liu et al., 2012) and PSB-2113 (PDB: 7PX4) (Claff et al., 2022), AB928 displaces 4 and 3 water molecules in the ligand binding pocket, respectively (Figure S5). These waters within the pocket are referred to as “unhappy water”, and their displacement by ligands is considered energetically favorable (Mason et al., 2013). It is worth noting that another clinical investigational drug, AZD4635, also extends toward the secondary pocket and displaces several waters within the pocket, despite having one of the smallest receptor-binding interfaces among typical antagonists (Figure 2D) (Borodovsky et al., 2020).

We propose potential binding models for AB928/2-118 on A_2B_R. AB928 may undergo a subtle rotation towards V250^6.51^ to occupy the space left by the L/V^6.51^ variation, and its pyridine ring moves slightly out of the secondary pocket and makes contacts with the basic residues at ECL3. In contrast, the pyridine ring in 2-118 may collide with TM6 during rotation due to its different linkage. While both AB928 and 2-118 can rotate around the C-C and C-N bonds between the pyridine and the triazole/pyridinone rings when dock to their pockets, the rotations in 2-118 are evidently more restricted because of its larger pyridinone ring (Figure 4D). Therefore, 2-118 may need to make further adjustments in other directions, potentially disrupting interactions in the orthosteric pocket (e.g., the hydrogen bond contributed by the cyano goup), this is probably the reason why, despite its similarity to AB928, its inhibitory potency is greatly compromised. These models on A_2B_R reveal both similar and distinct features compared to the binding modes observed in A_2A_R crystal structures, and effectively explain our mutagenesis data on A_2B_R.

While we were preparing our manuscript, Claff *et al*. reported crystal structures of A_2A_R–AB928 with thermostabilized mutations (Claff et al., 2023). The binding mode revealed in that structures is largely similar with our AB928-bound WT A_2A_R structure with only tiny vibrations. However, that study did not investigate into the ligand’s potential binding mode on A_2B_R, thus advanced little on the dual-antagonism mechanism on A_2A_R/A_2B_R.

In conclusion, the crystal structures of A_2A_R in complex with AB928/2-118, along with the cAMP assay and MD simulations performed on both A_2A_R and A_2B_R, provide evidence that each ligand adopts a unique binding model on A_2B_R, which potentially explain their different inhibitory efficacies between A_2B_R and A_2A_R. This study can be used as structural templates for future development of selective or dual inhibitors against A_2A_R/A_2B_R for the treatment of related cancers.

## MATERIALS AND METHODS

### A_2A_R construct design, expression and purification

Human A_2A_R (residues 2-316) was cloned into a modified pFastBac1 vector containing a hemagglutinin (HA) signal peptide and a FLAG tag at the N-terminus, 10×His-tag at the C-terminus. In order to facilitate protein crystallization, the ICL3 of A_2A_R (residues K209–A221) was replaced with bacteriophage T4 lysozyme (T4L). Recombinant baculovirus expressing A_2A_R was prepared using Bac-to-Bac system (Invitrogen). Spodoptera frugiperda 9 (Sf9) insect cells were cultured in ESF921 medium and infected by 1%(v/v) high-tilter baculoviruses at the density of 2–3×10^6^ cells/ml. 1 L of Sf9 cells expressing A_2A_R were harvested 60 hours post infection by centrifugation, flash-frozen in liquid nitrogen and stored at −80°C for purification.

Cell pellets were thawed and resuspended using dounce tissue grinder in a hypotonic buffer containing 10 mM HEPES pH 7.5, 10 mM MgCl_2_, 20 mM KCl and EDTA-free protease-inhibitor cocktail (Bimake) twice, followed by three washes of a high salt buffer containing 10 mM HEPES pH 7.5, 10 mM MgCl_2_, 20 mM KCl, 1 M NaCl with EDTA-free protease-inhibitor cocktail. The membrane was collected by centrifugation at 150,000 ×g during each procedure above, then resuspended by the hypotonic buffer described above with an addition of 4 mM theophylline (Sigma), 2.0 mg/ml iodoacetamide (Sigma) and EDTA-free protease-inhibitor cocktail. After a 30-min incubation at 4°C in the dark, the membranes were solubilized by incubating with an addition of 1% (w/v) n-dodecyl-β-D-maltoside (DDM, Anatrace) and 0.2% (w/v) cholesterol hemisuccinate (CHS, Sigma) for 3.5 h at 4°C. Insoluble materials were removed by centrifugation at 150,000 ×g and the supernatant was isolated, added with 0.8 ml pure TALON IMAC (Clontech) resin and 20 mM imidazole, left to rock gently at 4°C overnight. The resin was washed with 2×10 column volumes (CV) of wash buffer 1 (25 mM HEPES pH 7.5, 500 mM NaCl, 5% (v/v) glycerol, 0.05% DDM, 0.01% CHS, 30 mM imidazole and 20 μM AB928 or 2-118), followed by another 10 CV of wash buffer 2 (25 mM HEPES pH 7.5, 500 mM NaCl, 5% glycerol, 0.025% DDM, 0.005% CHS, 30 mM imidazole and 20 μM AB928 or 2-118) and then eluted with 3 CV of elution buffer (25 mM HEPES pH 7.5, 500 mM NaCl, 5% glycerol, 0.025% DDM, 0.005% CHS, 300 mM imidazole and 100 μM AB928 or 2-118). The elution was concentrated with an Amicon centrifugal ultrafiltration unit (Millipore) with 100 kDa molecular-weight cut-off (MWCO). The concentrated samples were checked using high-performance liquid chromatography (HPLC) and gel electrophoresis.

### Thermal-shift assay

N-[4-(7-diethylamino-4-methyl-3-coumarinyl)phenyl]maleimide (CPM, Sigma) was dissolved in DMSO at 4 mg/ml as stock solution and diluted 20 times using a buffer containing 25 mM HEPES, pH 7.5, 500 mM NaCl, 5% glycerol, 0.01% DDM, 0.002% CHS before use. 0.5–1.0 μg purified A_2A_R diluted in the same buffer above at a final volume of 49 μl was incubated with 1 μl diluted CPM. For A_2A_R prepared for thermal-shift assay, no antagonists were added during purification and each antagonist was added only to each sample at a final concentration of 80 μM. The thermal-shift assay was performed by a Cary Eclipse fluorescence spectrophotometer (Agilent) with an excitation wavelength set to 365 nm and an emission wavelength detected at 460 nm. All assays were performed over temperature ranging from 25 to 90°C. Data were processed with GraphPad Prism 8.0 (GraphPad Software) and nonlinear curve-fitting was performed using Boltzmann sigmoidal.

### Crystallization

Purified A_2A_R was co-crystallized with AB928 or 2-118 using lipid cubic phase (LCP) technology. Concentrated A_2A_R (>15mg/ml) was mixed with lipid [10% (w/w) cholesterol, 90% (w/w) monoolein] at a ratio of 2:3 (v:v, protein:lipid) in a custom 2×100 μl model 1700 Gastight glass syringe mixer (Hamilton) to prepare an LCP mixture. Then each well on a 96-well LCP sandwich plate (FAstal BioTech) was loaded with 50 nl of this mixture, followed by an overlay with 0.8 μl of different precipitant solution using NT8 automatic dispenser (Formulatrix), sealed with glass cover and stored at 18°C for crystal growth. Diffracting-quality A_2A_R–AB928 crystals were obtained in the condition containing 100 mM sodium cacodylate trihydrate, 120 mM ammonium tartrate dibasic and 32% PEG 400. Diffracting-quality A_2A_R–2-118 crystals were obtained in the condition containing 100 mM sodium cacodylate trihydrate, 200 mM sodium tartrate dibasic dihydrate and 30% PEG 400. Crystals were harvested using Dual-Thickness MicroMounts (MiTeGen) loops and kept in cryo-pucks stored in liquid nitrogen before diffraction study.

### Data collection and model building

X-ray diffraction data were collected on beamline 45XU with an automatic data collection program at the Japan synchrotron radiation SPring-8 facility with the 10μm beam with 0.1 s exposures and an oscillation of 0.1° per frame. For each crystal we collected totally 10° of diffraction data, and all data were then automatically processed with the program KAMO (Yamashita et al., 2018), and indexed, integrated and scaled using XDS (Kabsch, 2010). The structure was solved by molecular replacement with Phaser (McCoy et al., 2007) using the ZM241385-bound A_2A_R structure (PDB ID 3EML) as the search model. Resulting model refinement and rebuilding were performed using Phenix (Adams et al., 2010) and Coot (Emsley et al., 2010). Statistics are provided in Table S1. The 3D figures in this article were prepared with PyMOL Version 2.3 (PyMOL Molecular Graphics System, Schrödinger, LLC).

### Cell culture

HEK293 human embryonic kidney cells were purchased from the Cell Bank of the Chinese Academy of Sciences (Shanghai, China). HEK293 cells were maintained in DMEM medium (Gibco, USA) supplemented with 1% penicillin-streptomycin solution (Gibco, USA) and 10% fetal bovine serum (FBS, Gibco, USA) in a 37 °C humidified incubator with 5% CO_2_.

### GloSensor cAMP Assay

GloSensor cAMP Assay was conducted as described in our previously studies (He et al., 2022; Kumar et al., 2017). In brief, HEK293 cells in a 6-cm dish were transiently transfected with 1 μg of pGloSensor-22F cAMP plasmid (Promega, USA) and 1 μg of human wild-type or mutated A_2A_R or A_2B_R overexpression plasmid using polyethyleneglycol (6 μl, Yeasen, China). After 24-h incubation, transfected cells were harvested and re-seeded into 384-well white plates (Costar, USA) at a density of 20,000 cells per well in equilibration with CO_2_-independent medium (Gibco, USA) supplemented with 1% (v/v) GloSensor™ cAMP reagent (Promega, USA). Then, cells were pre-treated for 30 min with a series of concentrations of compounds and subsequently stimulated with NECA (MCE, USA). The bioluminescence intensity was acquired continuously for 30 min by a Cytation 5 imaging reader (BioTek, USA).

### MD simulations

For A_2A_R simulation systems, the simulations were initiated using the 2-118-bound and AB928-bound A_2A_R crystal structures with the T4L removed. For A_2B_R simulation systems, A_2B_R structures were modelled from 2-118-bound and AB928-bound A_2A_R crystal structures, and ligands were directly aligned to A_2B_R. Next, the CHARMM-GUI server (Wu et al., 2014) was used to insert them into POPC (palmitoyl-2-oleoyl-sn-glycero-3-phosphocholine) membrane. And TIP3P waters were added on the top and bottom of these simulation systems. In all, 0.15 mol/L NaCl ions and counterions were finally added to solvent. Neutral acetyl and methylamide groups were added to cap the N- and C-termini of protein chains, respectively. For each of these four simulation conditions, we performed 3 independent simulations in which initial atom velocities were assigned randomly and independently.

All MD simulations were performed using the GROMACS2020.2 package with the CHARMM36m forcefield (Huang et al., 2017). Parameters for ligands with high penalty scores were generated with CGenFF program (Vanommeslaeghe and MacKerell, 2012). Before the final production run of 500-ns simulations, 50,000 steps of energy minimization were performed for each system followed by equilibration in the NPT ensembles for 20-ns, with positional restraints (1,000 kJ mol^-1^ nm^-2^) placed on heavy atoms of protein and ligands. System temperature was maintained at 300 K using the v-rescale method with a coupling time of 0.1 ps and pressure was maintained at 1 bar using the Berendsen barostat with a coupling time of 1.0 ps and compressibility of 4.5 × 10^−5^ bar^-1^ with semi-isotropic coupling. A 2-fs timestep and LINCS constrained bond lengths were set during these simulations. Electrostatic interactions were computed using the particle mesh Ewald (PME) method with non-bonded interactions cut at 1.2 nm. The results of the MD simulations were analyzed by GROMACS tools.

## Acknowledgments

This work was supported by the National Key Research and Development Program of China (2018YFA0507001), the basic research program of Science and Technology Commission of Shanghai Municipality (21JC1402400), the National Natural Science Foundation of China (32171215, 81972828, 82172644, 82273857 and 81830083), and the National Key Scientific Infrastructure for Translational Medicine (Shanghai) (TMSK-2021-120). We thank the Instruments Sharing Platform of School of Life Sciences, East China Normal University. We also thank the support of ECNU Multifunctional Platform for Innovation (001) and National Super Computing Center in Zhengzhou for providing computational resources for this study. The synchrotron radiation experiments were performed at the BL45XU of Spring-8, Japan.

## Author Contributions

W.Y. optimized constructs, expressed and purified the complex proteins for crystallization studies, and edited the initial manuscript; Y.X. did the cAMP inhibition assay. Z.Q. did the MD simulations and analysis; C.Y. assisted molecular cloning and mutagenesis, X.Y. helped cell culture and expression; Z.C. synthesized the compounds for crystallization and assay; X. Q. and W.Y. guided the chemical synthesis; Y. H. oversaw the simulation and modelling; L.M. conceived the project and supported the research; L.W. guided and analysed the assay data, and edited the manuscript. S.G. supervised the project, determined the structures, analysed the data, and wrote the manuscript.

## Data availability

Atomic coordinates and structure factors for the A_2B_R–AB928 and A_2B_R–2-118 structures have been deposited in the Protein Data Bank with identification code 8JWZ and 8JWY, respectively.

## Supplementary Information for

Figures and tables

